# Detecting Selection in Low-Coverage High-Throughput Sequencing Data using Principal Component Analysis

**DOI:** 10.1101/2021.03.01.432540

**Authors:** Jonas Meisner, Anders Albrechtsen, Kristian Hanghøj

## Abstract

Identification of selection signatures between populations is often an important part of a population genetic study. Leveraging high-throughput DNA sequencing larger sample sizes of populations with similar ancestries has become increasingly common. This has led to the need of methods capable of identifying signals of selection in populations with a continuous cline of genetic differentiation. Individuals from continuous populations are inherently challenging to group into meaningful units which is why existing methods rely on principal components analysis for inference of the selection signals. These existing methods require called genotypes as input which is problematic for studies based on low-coverage sequencing data. Here, we present two selections statistics which we have implemented in the `PCAngsd` framework. These methods account for genotype uncertainty, opening for the opportunity to conduct selection scans in continuous populations from low and/or variable coverage sequencing data. To illustrate their use, we applied the methods to low-coverage sequencing data from human populations of East Asian and European ancestries and show that the implemented selection statistics can control the false positive rate and that they identify the same signatures of selection from low-coverage sequencing data as state-of-the-art software using high quality called genotypes. Moreover, we show that `PCAngsd` outperform selection statistics obtained from called genotypes from low-coverage sequencing data.

## 2 Introduction

Natural selection is the main driver of local adaptation. Instead of tracing the adaptive phenotypic trait, a “reverse ecology” approach is commonly applied [12], where the genetic variant encoding for a beneficial trait is first identified followed by the underlying mechanism of the adaptive phenotype. This enables mapping of the genetic architecture of phenotypic adaptability driven by natural selection ([6] for review on human populations). A common approach to identify candidates under selection is based on outliers in an empirical distribution of differentiation between two or more groups of predefined populations. In it simplest form, it finds the variants with the biggest difference in allele frequency between two predefined populations. One of many methods based on this notion is Population Branch Statistics [30], an estimator of genetic differentiation based on allelic changes estimated with the fixation index (*F_ST_*). It identifies candidate regions as strong deviations from an empirical distribution between a target population, a closely related sister population and an outgroup. However, homogeneous discrete groupings of the populations is required for many of these models, albeit exceptions exist [2].

The reduced expenses for whole genome DNA sequencing, thanks to advanced High-throughput DNA sequencing technologies, has facilitated larger sample sizes in population genetics studies in the recent years, including samples with similar genetic ancestry [4, 13, 29, 25, 19, 28]. Identifying signatures of selection in populations of similar genetic ancestry can results in arbitrary population assignments when using methodologies that require discrete groups of populations. This can lead to reduced power and increased false positive rates as allele frequencies are estimated from non-homogeneous populations. Instead of coercing samples into groups, an alternative approach is to account for the continuous cline of genetic differentiation in the selection analysis. Recent studies has shown that principal components analysis (PCA) of genetic data can detect signals of selection in continuous populations [14, 7]. Briefly, the idea is to use PCA to infer a weight for each variant which is scaled to reflect genetic drift. Variants with deviating statistics from the null distribution of what is expected under pure drift are candidates for selection. This approach has been applied to several dataset, including populations of humans [13, 7, 3], wheat [22], cod [26], turbots [18], and tiger mosquito [9].

Two commonly used software that accounts for continuous population differentiation when per-forming selection scans are `FastPCA` [7] and `pcadapt` [14, 23]. Both software use called genotypes as input to obtain the top *K* principal components (PCs) and variant weights through a truncated singular value decomposition (SVD) [10, 24]. However, they differ in their derived test statistics. `pcadapt` uses robust Mahalanobis distance [15] to evaluate all top *K* PCs for estimating *z*-scores, whereas FastPCA test normalized variant weights for each PC separately. Both test statistics follow *χ*^2^ distributions from which a *p*-value for each polymorphic site is obtained.

In this study, we extended the `FastPCA` [7] and `pcadapt` [14] selection statistics to account for genotype uncertainty by leveraging the PCs and variant weights estimated iteratively in the PCAngsd framework [17] using genotype likelihoods. This allows us to analyze low-coverage data and naturally impute missing data based on individual allele frequencies estimated from the top K inferred PCs. We apply the novel methods to populations of East Asian ancestry and European ancestry using the low-coverage data of the 1000 Genome Project [4] and demonstrate that we can identify known signatures of selection within these two ancestries. The candidates under selection were verified using the corresponding high quality genotype data from the 1000 Genome Project. The test statistics are implemented in the PCAngsd framework [17] that is available at https://github.com/rosemeis/pcangsd.

## 3 Materials and Methods

We assume that variable sites are diallelic and the major and minor allele are known such that genotypes are expected to follow a Binomial model. In low-coverage sequencing data, genotypes are unobserved and genotype likelihoods are therefore used instead to account for the uncertainty in sequencing process. We use the iterative procedure in `PCAngsd` [17] to estimate individual allele frequencies that can be seen as the underlying parameters in the Binomial sampling processes of the genotypes accounting for population structure. In the following, we will denote *N* as the number of individuals and *M* as the number of sites. We can then define the posterior genotype dosage as follows for individual *i* in site *j*

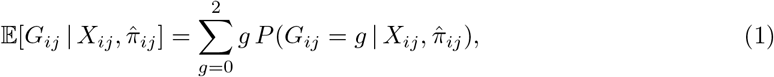

for *i* = 1,…, *N* and *j* = 1,…, *M*, where 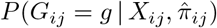 is the posterior genotype probability of genotype *g* with *X* being the observed sequencing data, and 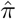 being the individual allele frequency. Details of deriving the posterior genotype from genotype likelihoods can be found in the supplementary material (Equation S1-S2). Missing data is imputed based on population structure based on the posterior genotype dosages. We standardize the dosage under the assumption of a Binomial model,

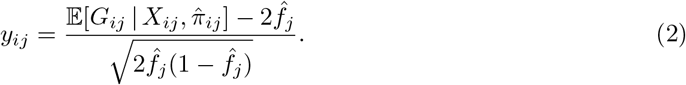

Here 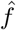 is the estimated allele frequency at site *j* based on all of the samples. We then perform truncated SVD [10] on the full standardized data matrix (*N* × *M*) to extract the top *K* principal components (PCs) that capture population structure in the dataset

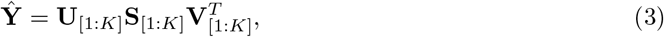

where **U**[_1:*K*_] represents the captured population structure of the individuals and **V**[_1:*K*_] represents the scaled variant weights, while **S**[_1:*K*_] is the diagonal matrix of singular values. This low-rank approximation along with the standardized matrix **Y** are all we need to estimate the two test statistics for low-coverage sequencing data.

## 3.1 FastPCA statistic

The selection statistic derived in Galinsky et al. (2016) [7], hereafter referred to as `FastPCA`, tries to detect selection by looking for variants that significantly differentiate from genetic drift along an axis of genetic variation. They define the selection statistics for the *k*-th principal component to be the properly normalized variant weights, using the properties of an eigenvector, such that they are standard normal distributed. The selection statistics are then defined as follows in our setting for genotype likelihood data

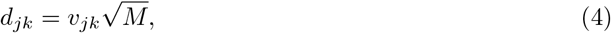

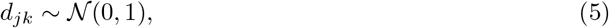

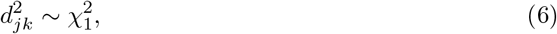

for *j* = 1,…, *M* and *k* = 1,…, *K*. *v_jk_* is the variant weight for the *k*th component at site *j*. The squared statistic will then follow a *χ*^2^-distribution with 1 degree of freedom. This statistic is implemented in the `PCAngsd` framework and referred to as `PCAngsd-S1`.

## 3.2 pcadapt statistic

The test statistic implemented in `pcadapt` [14] is based on a robust Mahalanobis distance of the standardized estimates in a multiple linear regression for each site. The regression model is defined as follows in our setting for genotype likelihood data

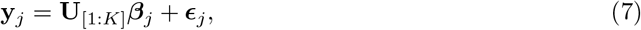

for *j* = 1,…, *M*, with *β_j_* being the regression coefficients, and *ϵ_j_*, the residual vector for site *j.* The coefficients are easily derived using the normal equation and properties of the previously computed truncated SVD (Equation 3), thus *β_j_* = **S**_[1:*K*]_ **V**_[*j*1:*K*]_. A *z*-score of the regression coefficients in site *j* are defined as

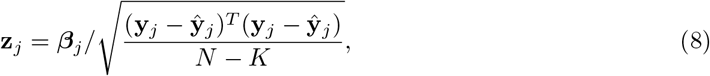

with 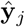 being the vector of low-rank approximations in site *j* (Equation 3). The test statistic is computed as a robust Mahalanobis distance of **z**_*j*_, where the squared distance will be 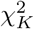 distributed as described in Luu et al. (2016) [14]. We use standardized expected genotypes *y_ij_* (Equation 2) for genotype likelihood data, hereafter referred to as `PCAngsd-S2`, instead of using known genotypes as `pcadapt`. Note, that we correct for inflation using the genomic inflation factor [5], inline with the recommendations [14], in all analysis based on the `pcadapt` or `PCAngsd-S2` statistics. See QQ-plot in Figure S1 and S2 for examples of the uncorrected `PCAngsd-S2` test statistic.

## 3.3 1000 Genomes Project data

We applied the two selection statistics implemented in the `PCAngsd` framework to the low-coverage data of the 1000 Genomes Project (phase 3). Specifically, we tested two sets of populations, one with East Asian ancestry with 400 unrelated individuals from four East Asian populations (CHB, CHS, CDX, and KHV), and one with European ancestry with 404 unrelated individuals from four European populations (CEU, GRB, IBS, and TSI). High quality genotype data is available for all the individuals analyzed. First, we calculated genotype likelihoods (GL) from the low-coverage data using `ANGSD` [8], restricting to polymorphic sites with a minor allele frequency of 5% in the high quality genotype data for each set of populations. In total 5.8 and 6 million polymorphic sites are retained in the Asian and European population sets, respectively. We used the GL data as input to `PCAngsd` to compute the two selection statistics (`PCAngsd-S1, PCAngsd-S2`) for the population sets. To verify the results obtained from the low-coverage data, we also analyzed the same variable sites from the high quality genotype (`HQG`) data using `PCAngsd, pcadapt` (default settings), and `FastPCA` (`fastmode:YES`, following [7]).

To compare the performance of `pcadapt` and `FastPCA` on low-coverage data, we called genotypes for the same variants described above from the low-coverage data using `bcftools` [11] and generated two data sets, one excluding all genotype calls with genotype quality < 20 (`CG standard`) and one including all called genotypes (`CG*`).

## 4 Results and Discussion

To test the performance of the two selection statistics (`PCAngsd-S1` and `PCAngsd-S2`), implemented in `PCAngsd`, on continuous genetic differentiation in low-coverage data sets, we used data from the 1000 Genomes Project [4]. We tested four populations with East Asian ancestry and four populations with European ancestry and identified known signatures of selection in both ancestries. We compared the results to `FastPCA` and `pcadapt` applied to `HQG` data and two data sets based on called genotypes from the low-coverage data, `CG standard` where all genotype calls with a genotype quality lower than 20 were excluded and `CG*` containing all called genotypes.

We applied the selection statistics to 400 individuals from four populations (CHB, CHS, KHV, CDX) with East Asian ancestry. First, we performed PCA on the GL data using PCAngsd [17] where we observed a continuous separation between the northern (CHB, CHS) and southern (KHV, CDX) populations on the first principal component (PC) (Figure 1). `FastPCA` and `pcadapt` obtained a similar pattern on the `HQG` data (Figure 1). PC2 obtained from `PCAngsd` and `pcadapt` separate the Vietnamese Kinh population (KHV) and Chinese Dai population (CDX) (Figure 1). When applied to the `CG standard` data, `FastPCA` and `pcadapt` could not recover the continuous separation on PC1. Instead we observe within population variance driven by the bias from genotype calling on low depth data when genotype quality filters are applied [20]. Therefore, `CG standard` data was not used for downstream selection scan comparisons. The PCA obtained from genotype data without quality filter `CG`* did not show the same problems and recovered the continuous separation on PC1 and was included in the following selection scan analyses 1.

**Figure 1:**
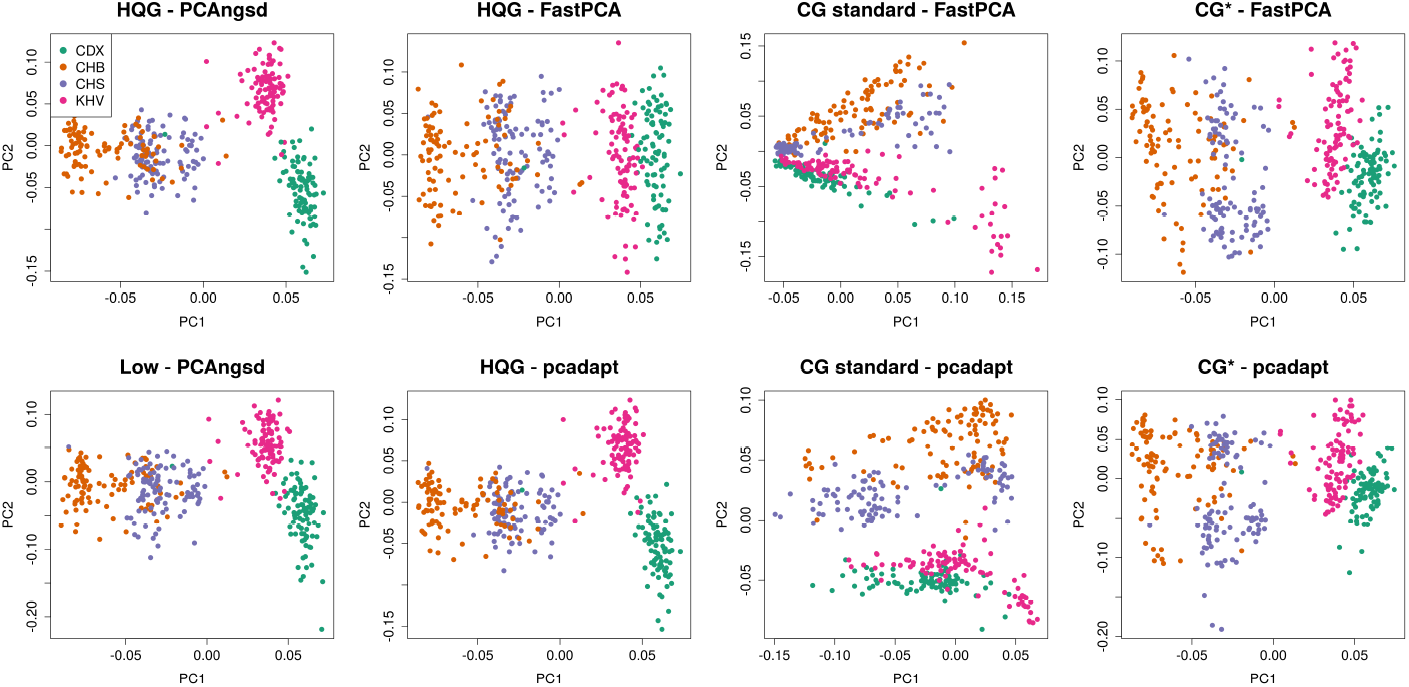
PC1 against PC2 of four East Asian populations obtained from `PCAngsd, FastPCA` and `pcadapt. HQG`: High quality genotype data, `Low`: Low-coverage data, `CG standard`: Called genotypes from low-coverage data with genotype quality threshold on 20, `CG*`: Called genotypes from low-coverage data.

We applied the test statistics on the variant weights inferred along the two PCs and scan for genomic regions with significant differentiation on the continuous north-to-south cline on PC1 and separation of KHV and CDX on PC2. We identify several candidates under selection along PC1 (Figure 2). After multiple testing correction using Bonferroni (*p*-value < 9 x 10^-9^, *α* = 0.05), we find significant signals of differentiation in variants overlapping *FADS2* (chr11), *IGH* cluster (chr14), *ABCC11* (chr16), and *LILRA3* (chr19). These signatures of selection have been described in previous studies of selection on continuous differentiation in Han Chinese populations [13, 3]. Interestingly, `PCAngsd` also identifies a genomic region overlapping *CR1* on the low coverage data, previously described by Chiang and colleagues [3] and the NIPT data [13]. We find a similar signal using the other software on the `HQG` although not significant. `FastPCA` and `pcadapt` find the same candidates with significant differentiation when applied to the `HQG` data.

**Figure 2:**
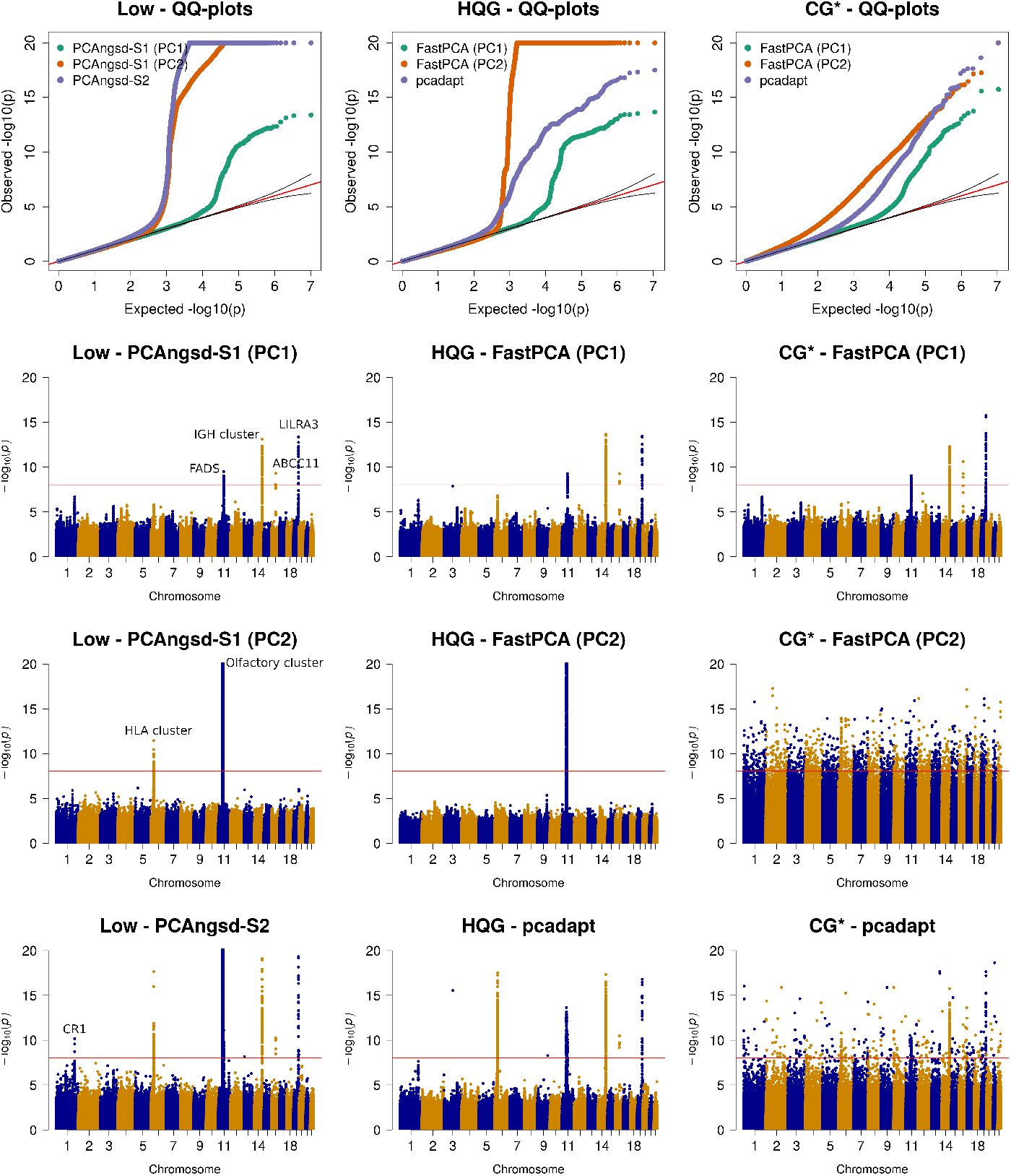
Selection scan of East Asian populations. QQ and Manhattan plots of the selection statistics from `PCAngsd`, `FastPCA` and `pcadapt` applied to the four East Asian populations obtained. Red horizontal line is the Bonferroni adjusted significance level. `PCAngsd-S2` and `pcadapt` has been corrected for genomic inflation. `HQG`: High quality genotype data, `Low`: Low-coverage data, `CG*`: Called genotypes from low-coverage data.

Both `PCAngsd` and `pcadapt` identify population structure on PC2 separating CDX and KHV and find the same two significant candidate regions: *HLA-cluster* (chr6) (also observed in [3]) and *Olfactory cluster* (chr11) (Figure 2). The variants overlapping the Olfactory cluster show strong LD pattern on both sides of the centromere, a challenging region to assemble potentially resulting in systematic biases, however, we do note that the pattern is present both on the `HQG` and low-coverage data (Figure 2 and S1). `PCAngsd-S2` and `pcadapt` identify a single significant variant on chr3 and chr9 in the `HQG` data. Following a test for Hardy-Weinberg equilibrium (HWE) accounting for population structure [16], we find that these two variants are the only top hits among selection candidates that significantly deviate from HWE (Table S1). This indicates genotype calling related biases as the variants are not candidates under selection in the low-coverage sequencing data.

When `FastPCA` and `pcadapt` are applied to the low depth data, `CG*`, not all of these signals are identified despite PC1 separating the four populations. We observed highly inflated statistics with significant false positive signals present genome-wide blurring the signals observed on the `HQG` data (Figure 2).

Similarly to the populations with East Asian ancestry, we also performed selection scans of 404 individuals from four populations (CEU, GBR, IBS, TSI) with European ancestry. We know from previous research that lactase persistence and skin and hair pigmentation distributions show a north-south cline within European populations [1, 21, 27], where the northern European populations have higher lactase persistence and lighter pigmentation than the southern European populations. We first performed PCA on the GL data using PCAngsd [17] (Figure 3) where we observed a continuous separation between the northern (CEU, GBR) and southern (TSI, IBS) populations on the first PC. `FastPCA` and `pcadapt` obtained a similar pattern on the `HQG` data (Figure 3). As for the East Asian scenario, `FastPCA` and `pcadapt` could not recover the continuous separation on PC1 on the `CG standard` data which was excluded from further analysis. The PCA obtained from `CG*` data set recovered the continuous separation on PC1 and was used in the following selection scan analyses 3.

**Figure 3:**
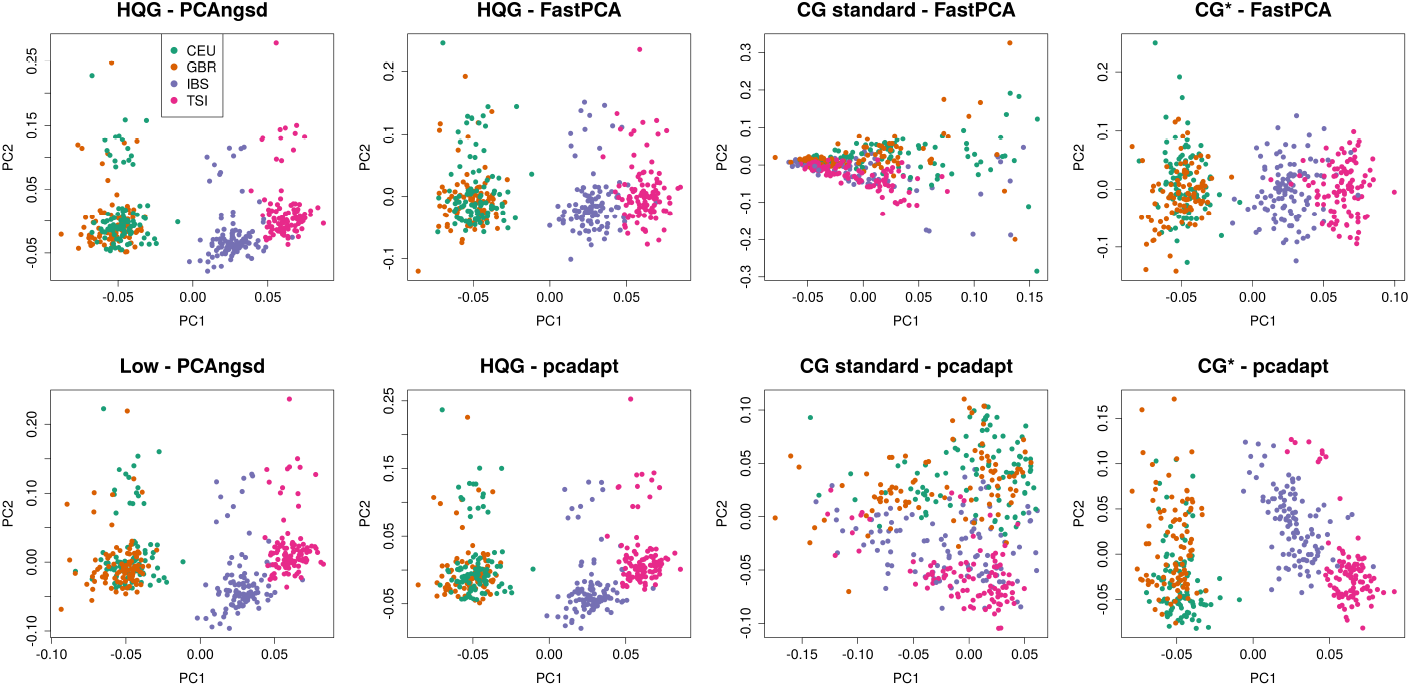
PC1 against PC2 of four European populations obtained from `PCAngsd`, `FastPCA` and `pcadapt`. `HQG`: High quality genotype data, `Low`: Low-coverage data, `CG*`: Called genotypes from low-coverage data.

Next, we calculated the selection statistics along PC1 that display a north-south cline in the European populations. We find that both `PCAngsd-S1` and `PCAngsd-S2` statistics behaves as expected under the null hypothesis for most sites(Figure 4). Similarly the statistics obtained from `FastPCA` and `pcadapt` follows the expectation, although, the latter required genomic inflation correction [5], on both `HQG` and `CG*`. After multiple testing correction, all software identify two genomic regions with significant genetic differentiation overlapping two gene clusters: *LCT/MCM6* (chr2) and *OCA2/HERC2* (chr15) (Figure 4). These results are inline with previous research on these populations [1, 21, 27].

**Figure 4:**
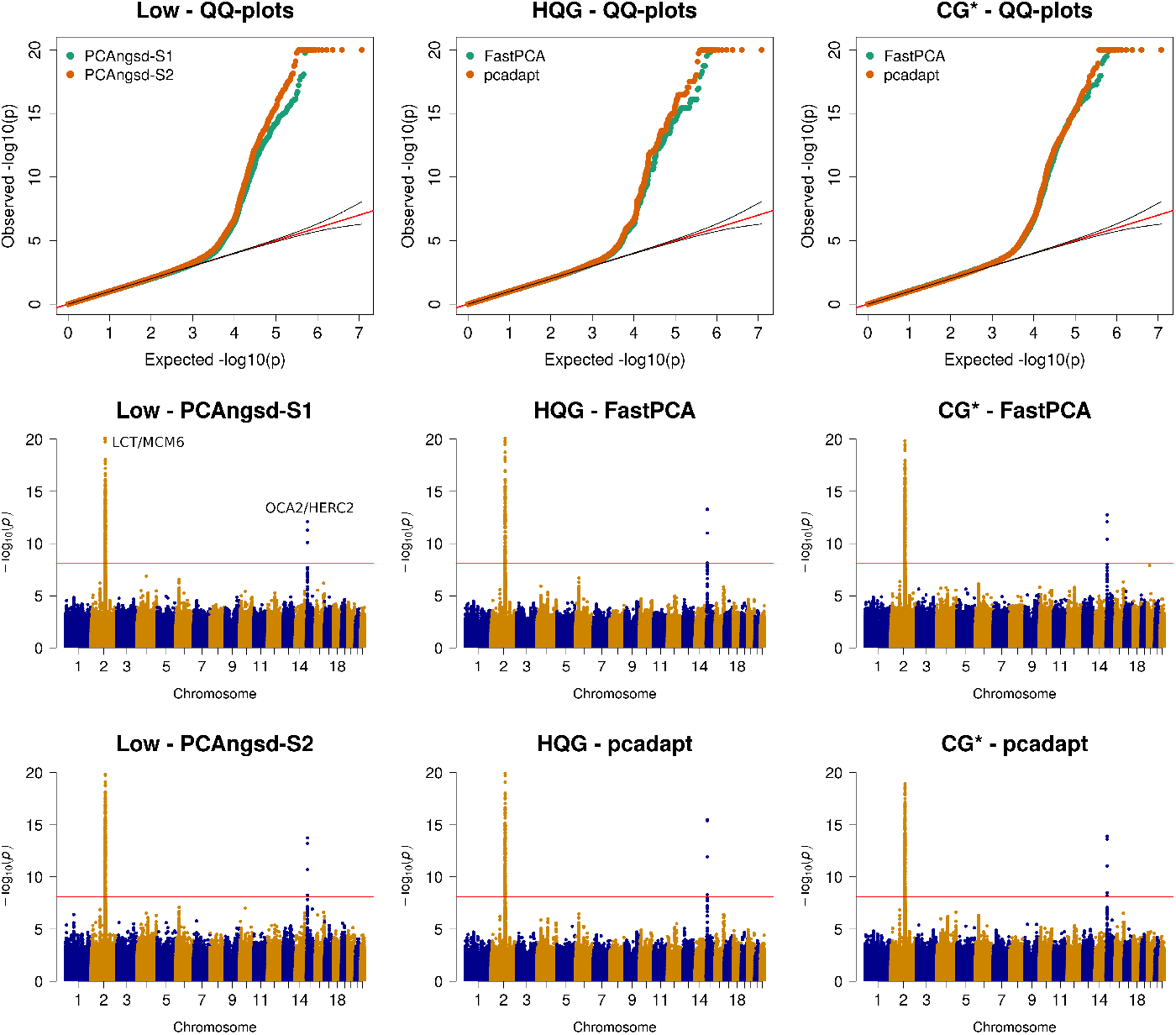
Selection scan of European populations. QQ and manhattan plots of the selection statistics from `PCAngsd`, `FastPCA` and `pcadapt` applied to the four European populations obtained. Red horizontal line is the Bonferroni adjusted significance level. `PCAngsd-S2` and `pcadapt` has been corrected for genomic inflation. `HQG`: High quality genotype data, `LOW`: Low-coverage data, `CG*`: Called genotypes from low-coverage data.

For genotype calling from low-coverage data uncertain genotype calls are often excluded by applying a genotype quality threshold. After applying a genotype quality threshold of 20 both `FastPCA` and `pcadapt` identify within population biases on the first PC (see Figure 1 and 3). However, as the second PC to some extent recover the population structure, we applied `FastPCA` and `pcadapt` to the standard genotype calls. In the selection scan of the East Asian populations `pcadapt` recovered the same candidates regions as the `HQG` data, whereas `FastPCA` identified many false positive regions both on PC1 and PC2 (Figure S3). For the European populations, we observe highly inflated statistics on both PCs and many false positive selection signatures were identified genome-wide by both software (Figure S4). From these observations, it is evident that genotype calling of low-coverage data requires ad-hoc filters for each test scenario. Similarly, in a recent low coverage study Chiang and colleagues also used extensive filters, including machine learning algorithms, to exclude outlier samples and variants prior to computing the selection statistics for the Han chinese population [3]. In contrast, we show that the `PCAngsd` framework consistently obtain well-behaving selection statistics in both scenarios from low-coverage data without the need for ad-hoc quality filters on either variant calls or sample selection.

A limitation of the PC-based selection scans are their capability of detecting selection in scenarios of non-continuous population structure. We show an example of this in Figure S5, where we have applied the three software to three populations with distinct ancestry (CEU, CHB, YRI). As also shown in the original study of `FastPCA` [7], it has low power in data sets with higher *F_ST_* between the populations, where we see deflated test statistics due to being inversely scaled with the inferred large eigenvalues of the corresponding tested PC for `PCAngsd-S1` and `FastPCA`. We see the opposite pattern for `PCAngsd-S2` and `pcadapt`, where the test statistics are very inflated, even after correction with genomic control, leading to many false positives. We reckon that *F_ST_* based selection scans are more appropriate in such scenarios with evident population clusters.

In conclusion, we have implemented two PC-based test statistics to perform selection scans in the `PCAngsd` (v.0.99) framework that performs iterative inference of population structure based on either GL or genotype data. This makes it possible to scan for selection genome-wide in data sets of low and/or variable coverage data sampled from genetically continuous populations. We show that the signatures of selection obtained from the low coverage in both the East Asian and European populations were on par with those from the high quality genotype data obtained from existing state-of-the-art software using called genotypes. The `PCAngsd` framework also reduces the need to rely on ad-hoc filters on SNP sites and/or samples. All obtained candidates for selection identified from the low-coverage data have been described in other studies targeting signatures of selection in European and East Asian ancestries. The `PCAngsd` framework is freely available at https://github.com/rosemeis/pcangsd.

## 5 Acknowledgements

The study was supported by the Lundbeck foundation.

## Supplementary Material

### Posterior expectation of the genotype

We are using the iterative algorithm in `PCAngsd` to estimate individual allele frequencies *π* [17]. With the assumption of Hardy-Weinberg proportions, we can derive the posterior genotype probability using the genotype likelihoods as follows for individual *i* in site *j*:

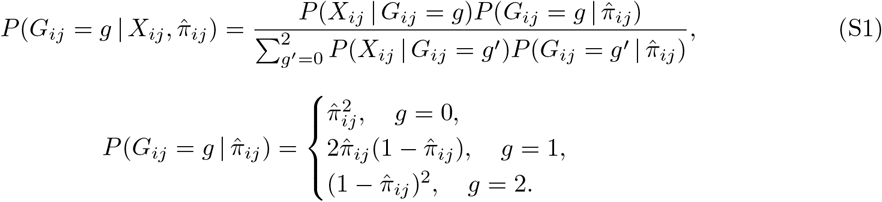

Here *g* is the genotype and *P*(*X* | *G* = *g*) is the genotype likelihood. The posterior expectation of the genotype is thus given by:

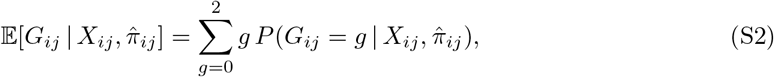

which we use in our selection statistics to account for uncertainty in the genotypes in low-coverage data.

### Supplementary figures

**Figure S1:**
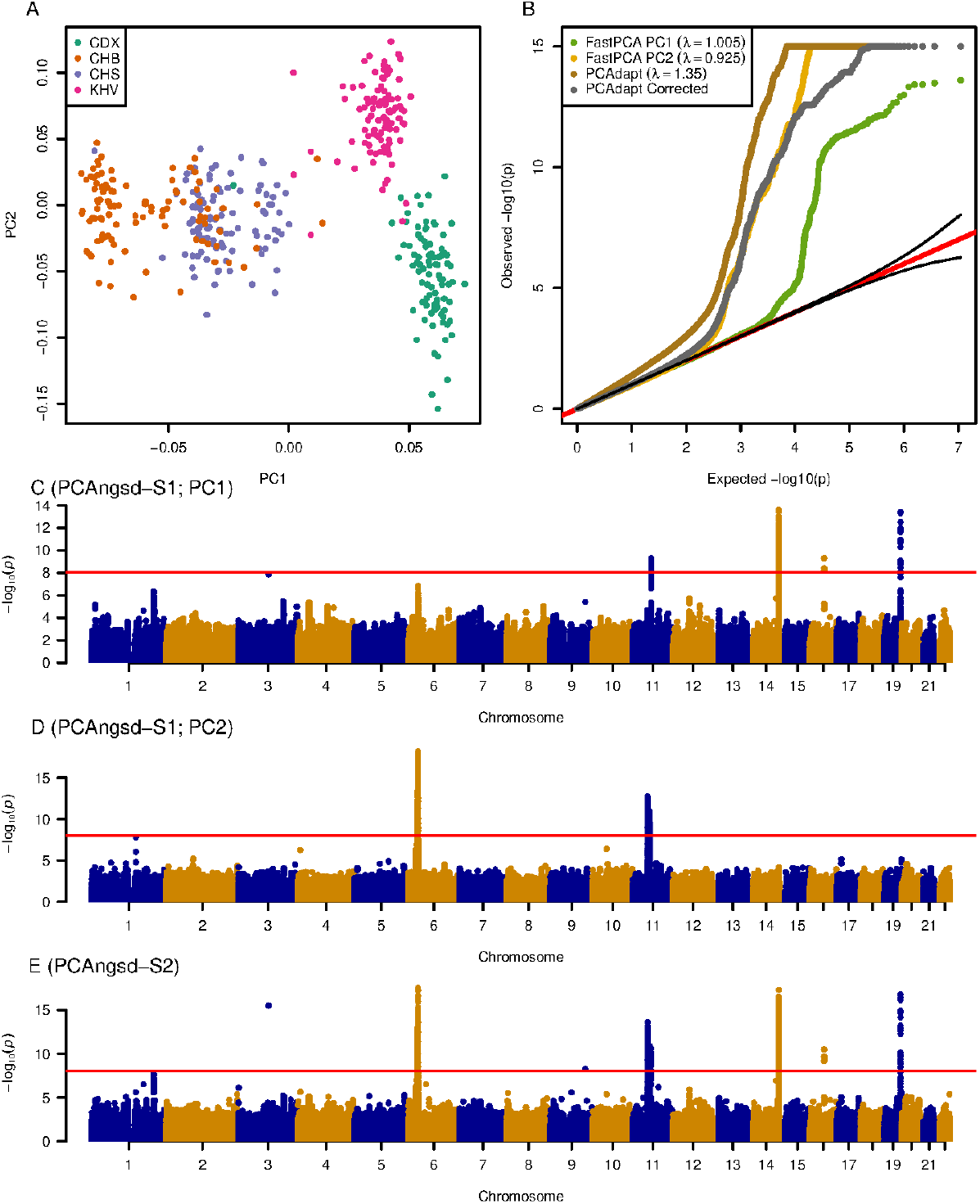
`PCAngsd` results on the high quality genotype dataset of the Asian populations in the 1000 Genomes Project. PCA plot of the four Asian populations showing the separation of Northern and Southern Asia on PC1 and PC2 separating KHV and CDX (A). QQ-plot of the test statistics, including `PCAngsd-S2` statistics before and after genomic inflation correction (B). Manhattan plot of the selection scan of PC1 (C) and PC2 (D) based on the `PCAngsd-S1` statistic and `PCAngsd-S2` (E) of both PCs. Manhattan plots from `PCAngsd-S2` has been corrected for genomic inflation. Red horizontal line is the Bonferroni adjusted significance level.

**Figure S2:**
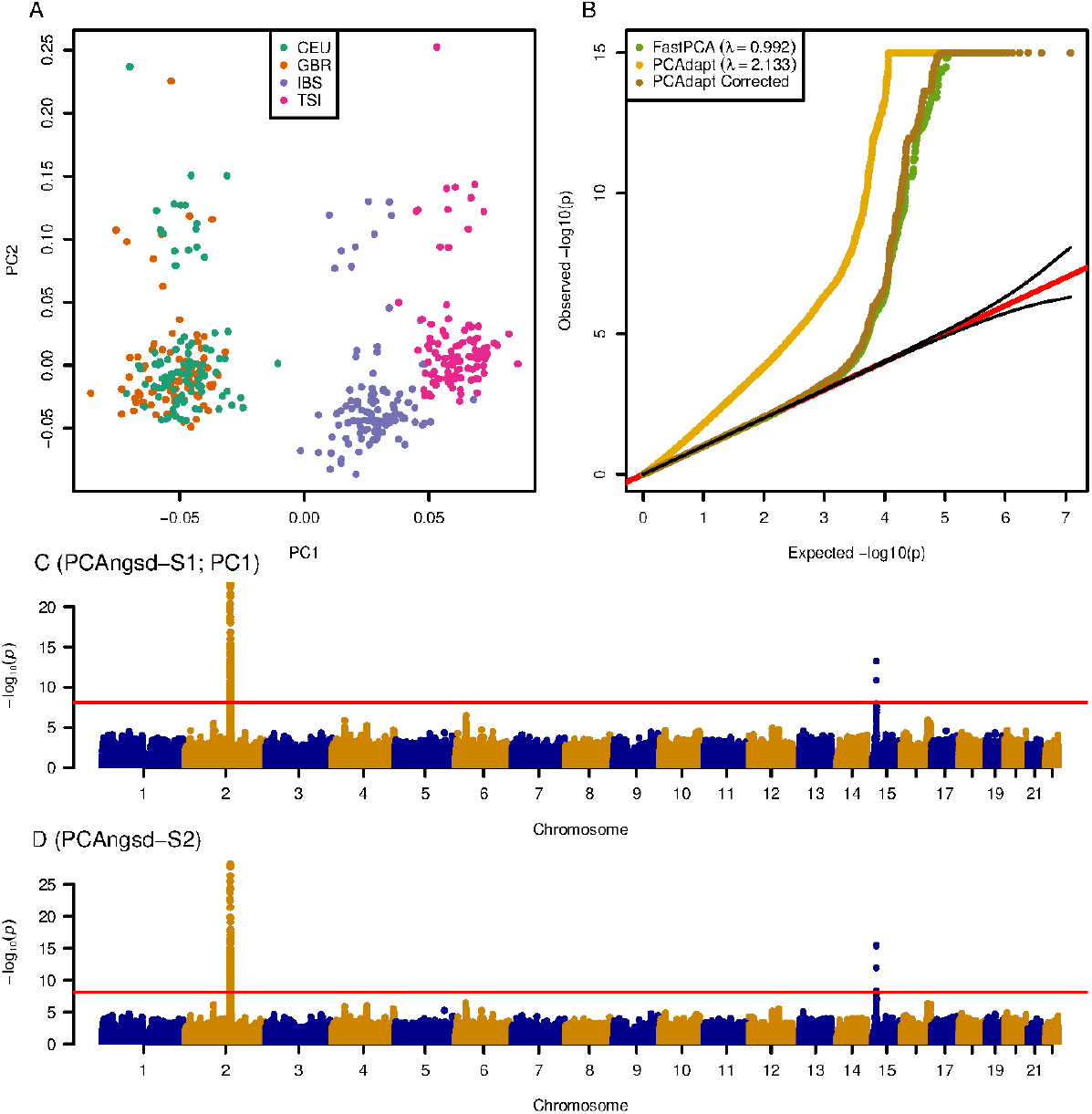
`PCAngsd` results on the high quality genotype dataset of the European populations in the 1000 Genomes Project. PCA plot of the four European populations showing the separation of Northern and Southern Europe on PC1 *(A).* QQ-plot of the test statistics, including `PCAngsd-S2` statistics before and after genomic inflation correction *B.* Manhattan plot of the selection scan based on the `PCAngsd-S1` *(C)* and `PCAngsd-S2` *(D)* test statistics along PC1. Manhattan plots from `PCAngsd-S2` has been corrected for genomic inflation. Red horizontal line is the Bonferroni adjusted significance level.

**Figure S3:**
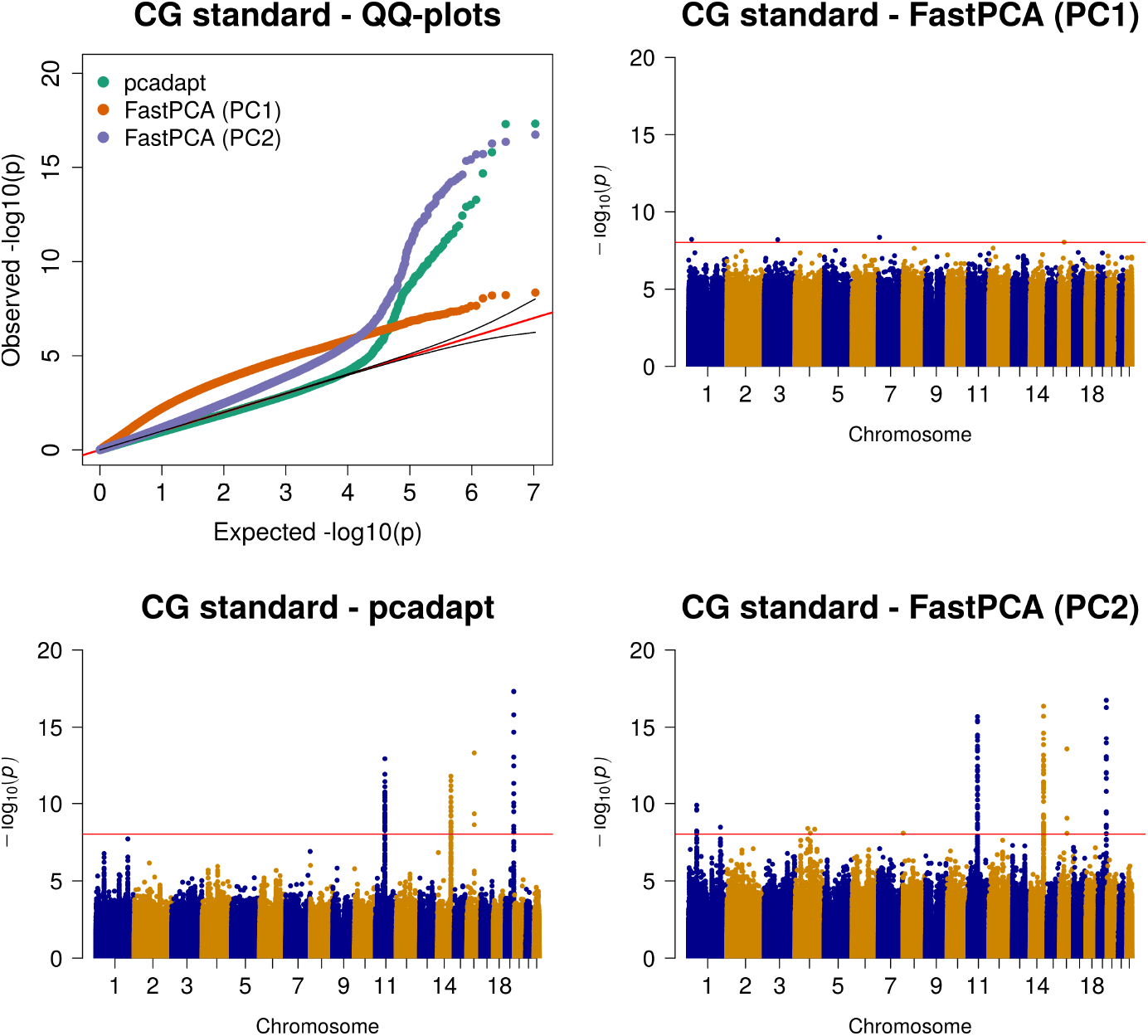
PC1 against PC2, QQ-plots and Manhattan plots of the selection statistics from `FastPCA` and `pcadapt` applied to the four East Asian populations obtained. Red horizontal line is the Bonferroni adjusted significance level. `pcadapt` has been corrected for genomic inflation. `CG standard`: Called genotypes from low-coverage data with a genotype quality threshold on 20.

**Figure S4:**
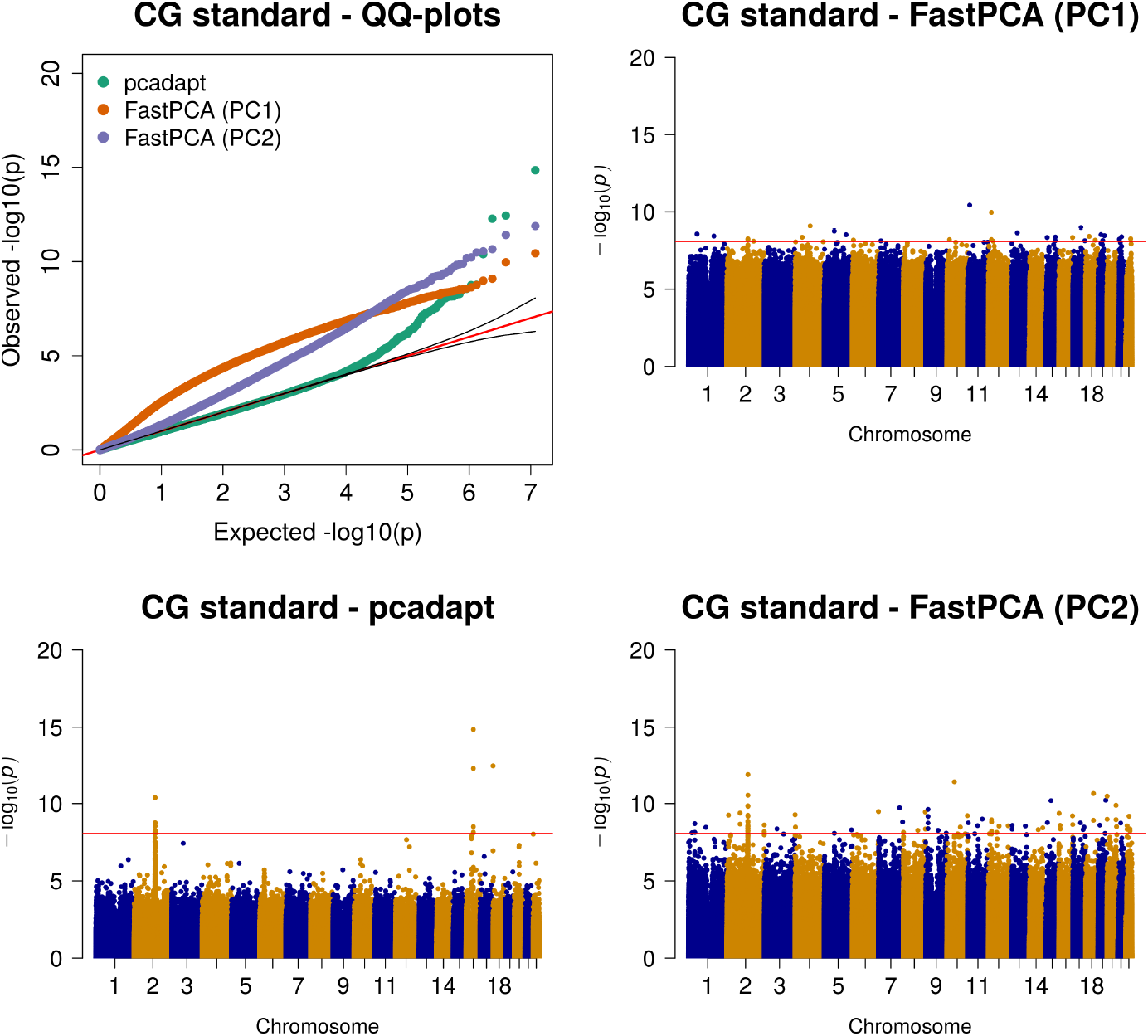
PC1 against PC2, QQ-plots and Manhattan plots of the selection statistics from `FastPCA` and `pcadapt` applied to the four European populations obtained. Red horizontal line is the Bonferroni adjusted significance level. `pcadapt` has been corrected for genomic inflation. `CG standard`: Called genotypes from low-coverage data with a genotype quality threshold on 20.

**Figure S5:**
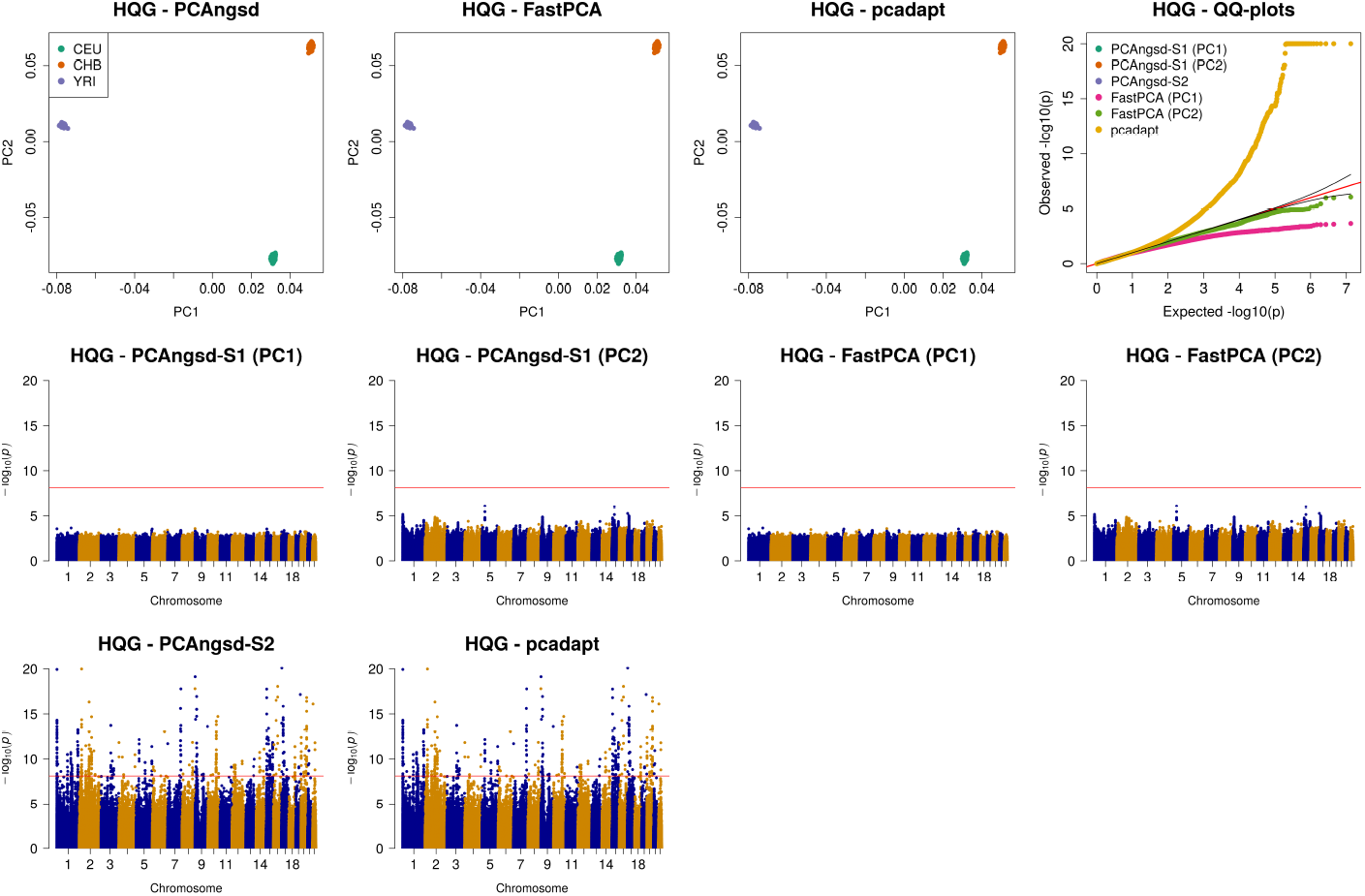
PC1 against PC2, QQ-plots and Manhattan plots of the selection statistics obtained from PCAngsd, `FastPCA` and `pcadapt` applied to a European (CEU), Asian (CHB), and African (AFR) population. Red horizontal line is the Bonferroni adjusted significance level. `pcadapt` has been corrected for genomic inflation. `HQG`: High quality genotype data.

**Table S1:** Hardy-Weinberg equilibrium test using `PCAngsd` on the `HQG` data from the four East Asian populations. The table only contains the significant top hits from the selection analyses. F: inbreeding coefficient.

| Chrom | ID | Position | A1 | A2 | F | $p$-value |
| --- | --- | --- | --- | --- | --- | --- |
| 3 | rs149768401 | 100365528 | C | G | -0.40 | $1.20 \times 10^{-12}$ |
| 6 | rs41542812 | 32629931 | G | C | 0.086 | 0.42 |
| 9 | rs115349067 | 117013044 | C | A | -0.26 | $1.26 \times 10^{-7}$ |
| 11 | rs7101761 | 49598178 | G | A | -0.068 | 0.071 |
| 11 | rs72643559 | 61620274 | C | T | -0.039 | 0.23 |
| 14 | rs1071803 | 106209119 | T | C | 0.022 | 1 |
| 16 | rs17822931 | 48258198 | C | T | -0.015 | 0.64 |
| 19 | rs434124 | 54809336 | C | G | -0.011 | 1 |

